# Distributed Visual Category Processing Across Medial Superior Temporal and Lateral Intraparietal Cortices

**DOI:** 10.1101/2020.08.25.266791

**Authors:** Yang Zhou, Krithika Mohan, David J. Freedman

**Author notes:** Corresponding author: David J. Freedman. Lead contact: David J. Freedman, Ph.D., Department of Neurobiology, The University of Chicago, Chicago, IL 60637, USA.

## Abstract

Categorization is an essential cognitive and perceptual process for recognition and decision making. The posterior parietal cortex (PPC), particularly the lateral intraparietal (LIP) area has been suggested to transform visual feature encoding into cognitive or abstract category representations. By contrast, areas closer to sensory input, such as the middle temporal (MT) area, encode stimulus features but not more abstract categorical information during categorization tasks. Here, we compare the contributions of PPC subregions in category computation by recording neuronal activity in the medial superior temporal (MST) and LIP areas during a categorization task. MST is a core motion processing area interconnected with MT, and often considered an intermediate processing stage between MT and LIP. Here we show that MST shows robust decision-correlated category encoding and working memory encoding similar to LIP, suggesting that MST plays a substantial role in cognitive computation, extending beyond its widely recognized role in visual motion processing.

## Introduction

Assigning incoming sensory stimuli into behaviorally relevant categories is essential for recognizing the significance of sensory information and generating task-appropriate behavioral responses. Previous studies, particularly those based on delayed match to category paradigms (DMC)^1,2^, have shown that several cortical areas, including the prefrontal cortex (PFC)^2-6^, posterior parietal cortex (PPC) ^1,3,7,8^, and inferior temporal (IT) cortex^9,10^, are involved in the visual categorization process. Recently, LIP (a subdivision of PPC) was shown to play a causal role in perceptual and categorical decisions about visual motion stimuli^11^. LIP also shows categorical encoding that is stronger, shorter in latency, and more strongly decision correlated than PFC activity^3^. Upstream visual motion processing areas, such as the middle temporal area (MT), show strong direction encoding, but not abstract categorical encoding, of visual motion during the same visual motion categorization task^1^. How motion direction encoding in MT is transformed into more cognitive categorical encoding in downstream areas, such as LIP, remains unclear. One possibility is that this transformation is achieved by flexible readout of MT activity by LIP. Alternatively, other brain areas may play a role in mediating this transformation. Here we address this question by directly comparing the roles of LIP and MST, an important parietal motion processing area that is reciprocally connected with both LIP and MT^12,13^, in visual-motion categorization in order to understand how sensory encoding of visual motion is transformed into flexible, learning-dependent, and task-related categorical representations.

MST has been identified as an important motion processing area within the dorsal visual pathway. MST is involved in the perception of both 2D and 3D visual motion patterns, and MST neurons typically have large receptive fields with responses that are selective for both simple motion stimuli as well as complex motion patterns such as “optic flow” stimuli as are generated by one’s own motion through the visual world^14-21^. It has also been suggested that MST is involved in transforming spatial information between different reference frames during self-motion^22,23^. Moreover, dorsal MST (MSTd) also integrates visual and vestibular signals^24-28^. However, MST’s contributions to cognitive functions have been less studied compared to other PPC areas such as LIP and 7a^22,29,30^. One recent study found striking motion-direction selectivity in MST during the delay-period of a delayed match to sample task^31^, whereas obvious selective delay period activity was not observed in MT spiking activity in that study, or in recordings from MT during a visual motion categorization task^1^. Several studies which compared neural activity between MST and LIP did find encoding of extraretinal or task-related factors in MST, but that encoding was typically weak in comparison with cognitive encoding in LIP^29,30,32^.

Here we directly compared neural activity between MST and LIP while monkeys performed a visual motion DMC task in which they needed to categorize a sample stimulus, maintain sample-category information in short-term memory, and determine whether a subsequent test stimulus was a categorical match or non-match to the sample. We found that MST neurons showed significant motion category encoding during stimulus presentation and memory delay periods of the DMC task, qualitatively similar to that observed in LIP. Also similar to LIP, MST category encoding was correlated with monkeys’ trial-by-trial categorical decisions, revealed by comparing category selectivity on correct vs. error trials. However, our analysis suggests that LIP is more closely involved in the categorical-decision process compared to MST, as decision-correlated category encoding in MST tended to be longer-latency and weaker than in LIP in the period immediately following stimulus presentation. Instead, category encoding in MST peaked later in the trial, around the time of transition from the sample to memory delay. Furthermore, both spiking and local field potential (LFP) neural activity in MST showed substantial categorical encoding during the working memory period, and that encoding correlated with the monkeys’ trial-by-trial categorical choices. In summary, our results show that MST encodes abstract and task-related factors such as categorical decisions and working memory, going beyond its traditionally recognized role in visual motion processing. This also gives insight into the functional roles of hierarchically interconnected PPC subregions in perceptual and cognitive functions.

## Results

### Task and behavioral performance

We trained two monkeys to perform a visual-motion DMC task, in which they needed to decide whether two sequentially presented motion stimuli were a categorical match or non-match, which they reported by releasing or holding a manual touch-bar respectively (**Fig. 1a**). To solve the task, monkeys needed to first categorize the sample stimulus and remember it during the delay period in order to compare it to the category of the upcoming test stimulus. The stimulus set consisted of ten directions of 100% coherence random-dot movies that were assigned to two learned categories (five motion directions per category) by an arbitrary category boundary (**Fig. 1b**). There were two near-boundary directions in each category which were 10° from the boundary, while the other 6 directions were evenly spaced (60°). Both monkeys performed the DMC task with high accuracy during both MST and LIP recording sessions (**Fig. 1c**, monkey M: MST = 90.9 %± 4.0%, LIP = 87.6% ± 4.3%; monkey Q: MST = 89.5 % ± 2.7%, LIP = 86.7% ± 3.8%), and greater than 70% correct for the near-boundary directions. Monkeys’ accuracies were significantly greater for sample-directions that were farther from (≥30 degree) the boundary as compared with the near-boundary directions for both MST and LIP recording sessions (near vs. far: (monkey M, MST: 79.4% vs. 98.9%, p = 1.15e-35, df = 81, tstat = -21.8; LIP: 77.7% vs. 94.2%, p = 1.034e-34, df = 61, tstat = - 26.0); (monkey Q, MST: 80.0% vs. 95.9%, p = 3.13e-32, df = 52, tstat = -26.96; LIP: 75.2% vs. 94.4%, p = 3.26e-28, df = 37, tstat = -31.3); paired t test).

**Fig. 1.**
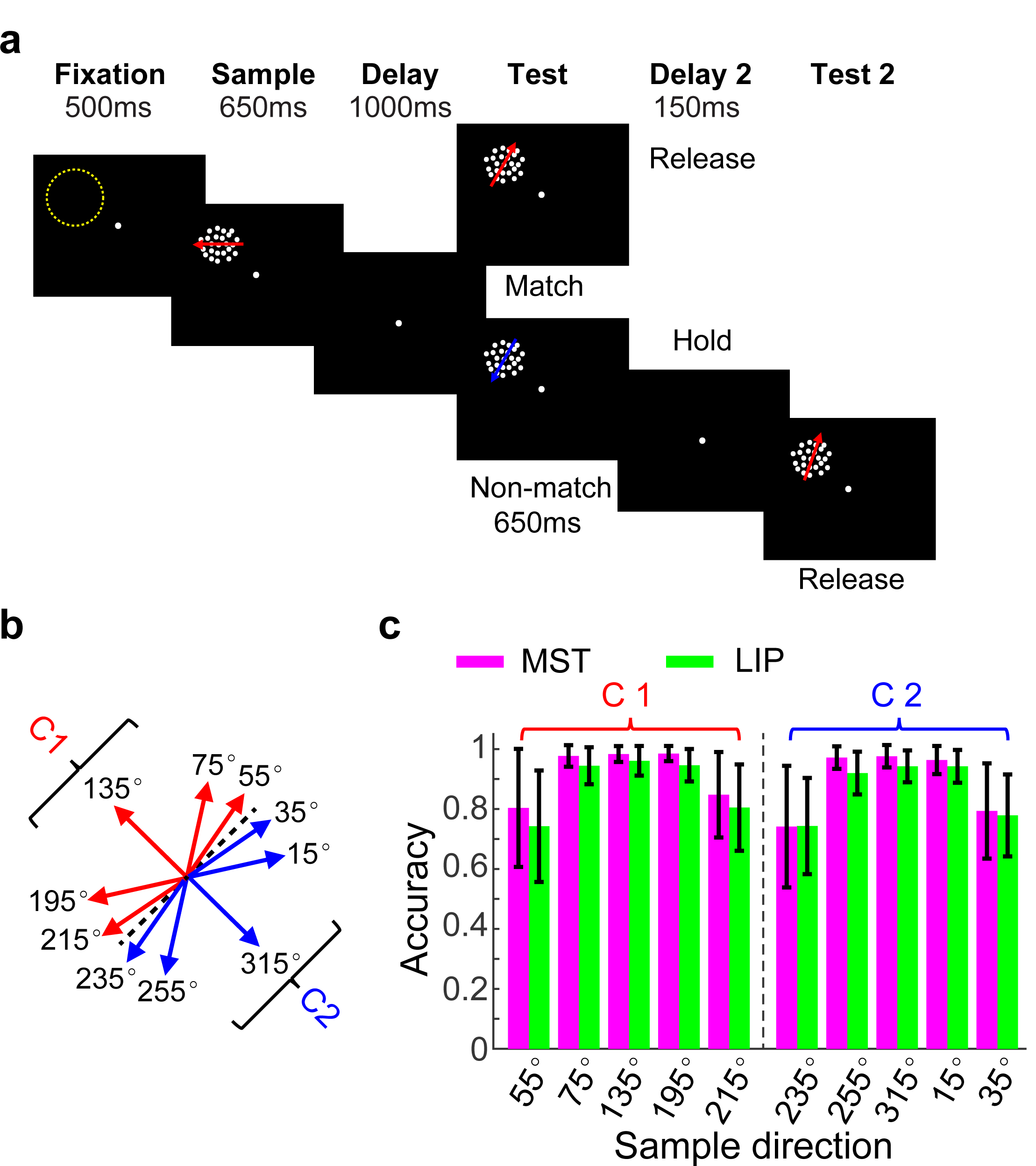
Task, behavioral performance and recording positions. **a**. Sequence of the DMC task. Monkeys needed to either release a touch-bar when the categories of sample and test stimuli matched, or hold the bar and wait for the second test stimulus when they did not match. The yellow dashed circle indicates the position of a neuron’s receptive field. **b**. Monkeys needed to group ten motion directions into two categories (corresponding to the red and blue arrows) separated by a learned category boundary (black dashed line). **c**. Two monkeys’ averaged performance (accuracy) for the ten sample direction conditions during MST and LIP recording sessions are shown separately. Error bars denote the ±STD across sessions.

### Category selectivity in MST and LIP

We recorded neuronal spiking activity and LFP signals from MST and LIP (targeting one brain area per session) while monkeys performed the DMC task (**Fig. 1d**). In total, we isolated 648 and 361 single neurons from MST (Monkey M: 140, Monkey Q: 508) and LIP (Monkey M: 89, Monkey Q: 272), respectively. 571 MST neurons (Monkey M: 105, Monkey Q: 466) and 326 LIP neurons (Monkey M: 78, Monkey Q: 248) showed task related responses in the DMC task (see criterion in *Methods*). A large proportion of neurons recorded from both areas showed significant category encoding in the DMC task (MST: 407 of 571, Monkey M: 68, Monkey Q: 339, LIP: 203 of 326, Monkey M: 58, Monkey Q: 145), see criterion in *Methods*). **Fig. 2a,b** shows two example MST neurons. The first one (**Fig. 2a**) showed binary-like responses to the sample category during the sample, delay and test periods of the DMC task; while the second neuron (**Fig. 2b**) showed significant category encoding mainly during the late sample and delay periods. **Fig. 2c,d** shows two example LIP neurons which encoded the sample categories during the sample, delay, and test periods. We also identified 68 MST neurons and 56 LIP neurons which were direction-tuned, but did not show any obvious sample category selectivity (see criterion in *Methods*, example neurons in **Extended Data Fig.1**).

**Fig. 2.**
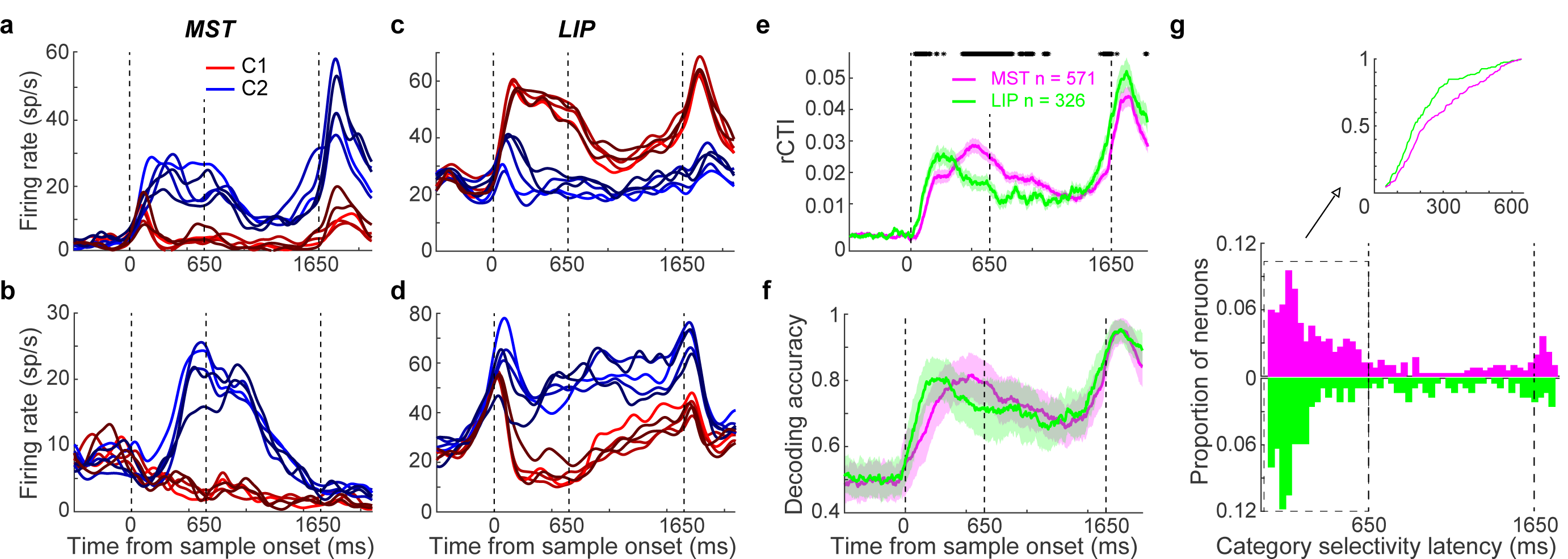
Sample category selectivity in MST and LIP during the DMC task. **a-b** Two example neurons from MST showed significant sample category selectivity. Average activity for each sample direction is plotted as a function of time. Different colors represent different sample categories, and different shades indicate the angular distance from the boundary. The first and second dashed vertical lines denote the time for sample stimulus onset and offset, respectively, while the third dashed vertical line indicates the time of test stimulus onset. **c-d**. Two example neurons from LIP are shown in the same format as a,b. **e-f**. Population level sample category selectivity in MST (pink) and LIP (green). **e**. A category tuning index (rCTI) shows the magnitude and time course of sample category selectivity. Shaded area denotes ±SEM. The black stars indicate the time points for which there was a significant difference between MST and LIP (unpaired t test, p < 0.05). **f**. Time course of sample category classification accuracy in MST (pink) and LIP (green) is measured using an SVM classifier. Shaded area denotes ±STD. g. The distributions of category selectivity latencies for category-selective neurons in MST and LIP. The above inset shows the cumulative distributions of category selectivity latencies only for neurons that were category-selective during the sample period.

To characterize the strength and time-course of neuronal sample category encoding across the MST and LIP populations, we calculated an ROC-based category tuning index (rCTI) for each single neuron recorded from both areas. As used previously^8^, the rCTI index quantifies each neuron’s category selectivity by comparing neuronal discriminability between pairs of directions in the same vs different categories (see *Methods*). rCTI values can range from -0.5 to +0.5 with more positive values indicating stronger category selectivity. As shown in **Fig. 2e**, the averaged sample rCTI values of both MST and LIP neurons are significantly shifted toward positive values shortly after sample onset, and maintained significantly positive values during the delay and test periods. This indicates that both MST and LIP showed significant category encoding during all periods of the DMC task. Comparing the two areas, LIP showed greater rCTI values during the early sample period (50 to 350ms after sample onset, p = 0.027, df = 905, tstat = -2.22, unpaired t test), while MST showed significantly greater category encoding during the late sample and early delay periods (−300 to 300ms relative to sample offset, p = 0.003, df = 905, tstat = -2.97, unpaired t test). Meanwhile, both MST and LIP neurons showed significant encoding of test stimulus category during the test period (**Extended Data Fig. 2**).

The latency of category selectivity for each neuron in both areas was determined by assessing the time-course of the rCTI (see *Methods*). This revealed that the distributions of category-selectivity latencies (across all the category-selective neurons) were similar between two areas (**Fig. 2g**, p = 0.144, zval = -1.46, ranksum = 75108, Wilcoxon test). To examine whether either brain area plays a leading role in categorizing sample stimuli, we compared the timing of category selectivity among those neurons that were category-selective during the sample period. We found that category encoding tended to emerge with a shorter latency in LIP than MST (**Fig. 2g**, p = 4.83e-04, zval = -3.49, ranksum = 3.40e+04, Wilcoxon test): a significantly higher proportion of neurons in LIP than MST were category-selective in the early sample period (50-350 ms following sample onset, p = 0.036, chi2stat = 4.38, chi-square test), and this trend was reversed during the late sample period (351-650 ms after sample onset, p = 1.16e-04, chi2stat = 14.85, chi-square test).

We next assessed category information at the population level using linear classifiers (support vector machine (SVM)) trained on neuronal pseudopopulations to decode the sample category on a trial-by-trial basis (see *Methods*). To mitigate the contribution of the category-independent direction tuning to category decoding performance, we trained and tested the classifier using different groups of directions in each category with similar angular distances (**Extended Data Fig. 3a**) as in previous studies from our group^4,8^. As shown in **Fig. 2f**, we found significant sample-category decoding in both MST and LIP in the sample, delay, and test periods. Interestingly, category encoding in MST emerged more gradually than LIP, reaching its initial peak during the late sample period. By contrast, LIP category decoding showed an initial peak during the early sample period. To assess category-independent motion direction encoding (i.e. the ability to discriminate between directions in the same category) in both MST and LIP, we trained linear direction classifiers (SVM) to decode the sample direction within each category (see *Methods* and **Extended Data Fig.3b**). This revealed substantial direction decoding in both MST and LIP primarily during the sample period (**Extended Data Fig. 4**).

**Fig. 3.**
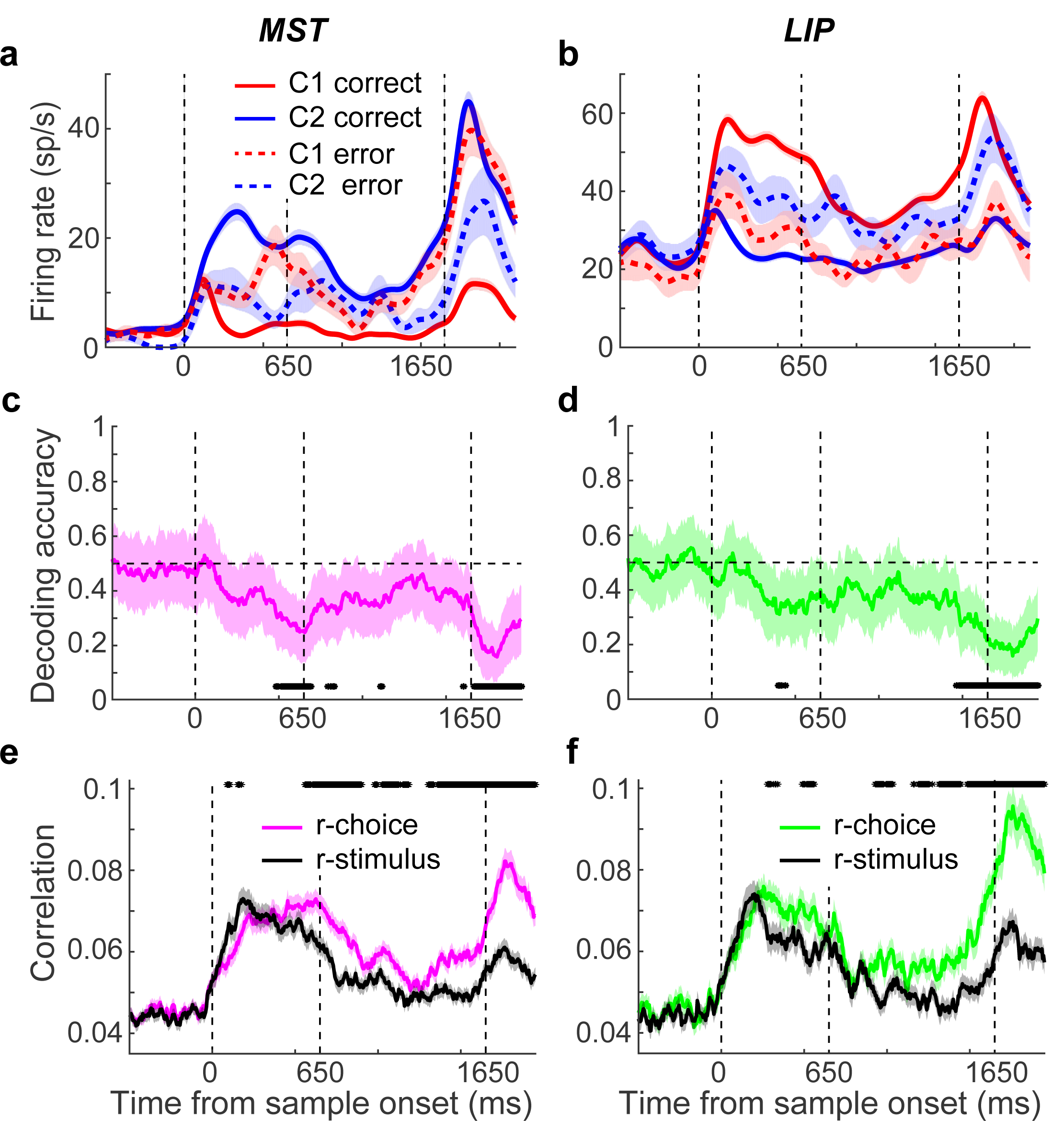
Sample category encoding in MST and LIP correlated with monkeys’ categorical decisions. **a**. An example MST neuron’s activity on correct (solid) and error (dashed) trials. Different colors represent different sample categories, and shaded areas denote ±SEM. **b**. An example neuron from LIP. **c-d**. The magnitude and time course of sample category selectivity on error trials in MST (c) and LIP (d) were determined by category SVM classifier. The shaded area denotes ±STD. The black stars indicate the time points for which the decoding performance was significantly lower than chance level (bootstrap, p < 0.05). **e-f**. Stimulus- and choice-related components of category selectivity in MST (e) and LIP (f) were determined by a partial correlation analysis. The values of r-stimulus (the partial correlation between neuronal activity and stimulus category, given the monkeys’ choices) and r-choice (the partial correlation between neuronal activity and monkeys’ choice, given the stimulus category) are plotted across time. The black stars indicate the time points for which there was a significant difference between r-choice and r-stimulus (paired t test, p < 0.05).

**Fig. 4.**
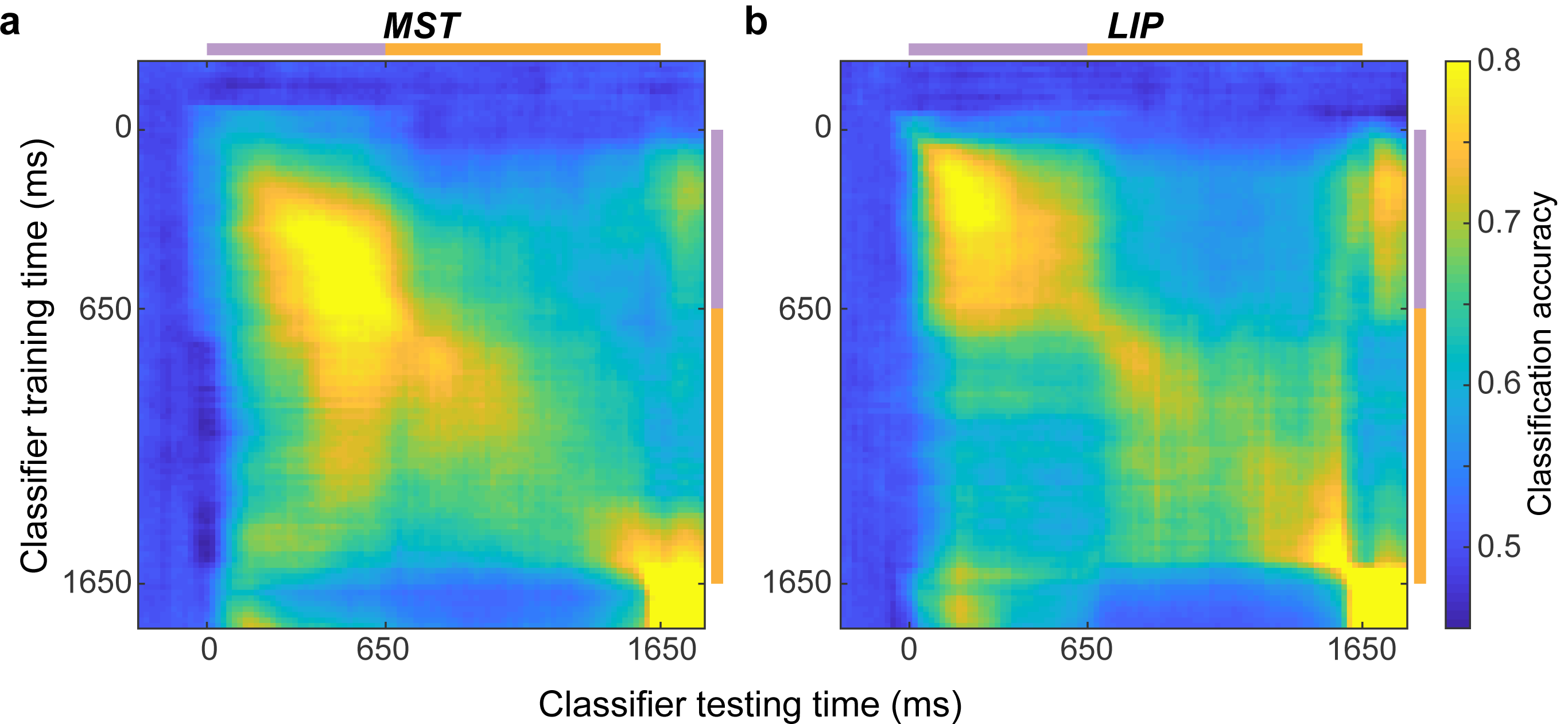
Stability of category selectivity in MST and LIP. **a**. The stability of MST sample category encoding was determined by training the classifier at one time point (y axis) and testing at other time points (x axis). Classification accuracy is indicated by color at each x-y coordinate. The purple and orange bars indicate the timing of the sample and delay periods, respectively. **b**. The stability of sample category encoding in LIP.

We were interested in the relationship between category encoding in both cortical areas and task difficulty. We assessed it by comparing category selectivity between the trials in which the sample direction was near vs. far from the category boundary. Because the more difficult (near-boundary) directions were spaced differently than the directions closer to the center of the categories, we could not use the rCTI or category classifier to directly compare the category selectivity for directions that were close vs. far from the category boundary. To address this, we used an unbiased fraction explained variance (FEV) analysis (see *Methods*) to quantify the selectivity among the five opposite direction pairs (180 degrees apart). We grouped the five direction pairs into more-difficult and easier groups (based on distance from the boundary) and then averaged the selectivity of the direction pairs belonging to each group. We found that sample category selectivity was significantly greater for more difficult trials than for easier trials in both MST and LIP during the delay period of the DMC task (MST: p = 5.99e-14, df = 566, tstat = 7.70; LIP: p = 1.33e-6, df = 317, tstat = 4.93; paired t test, **Extended Data Fig. 5**). This is consistent with a previous rodent study showing stronger category selectivity for auditory stimuli near the category boundary^33^, and suggests that category encoding in both MST and LIP was correlated with the levels of task difficulty.

**Fig. 5.**
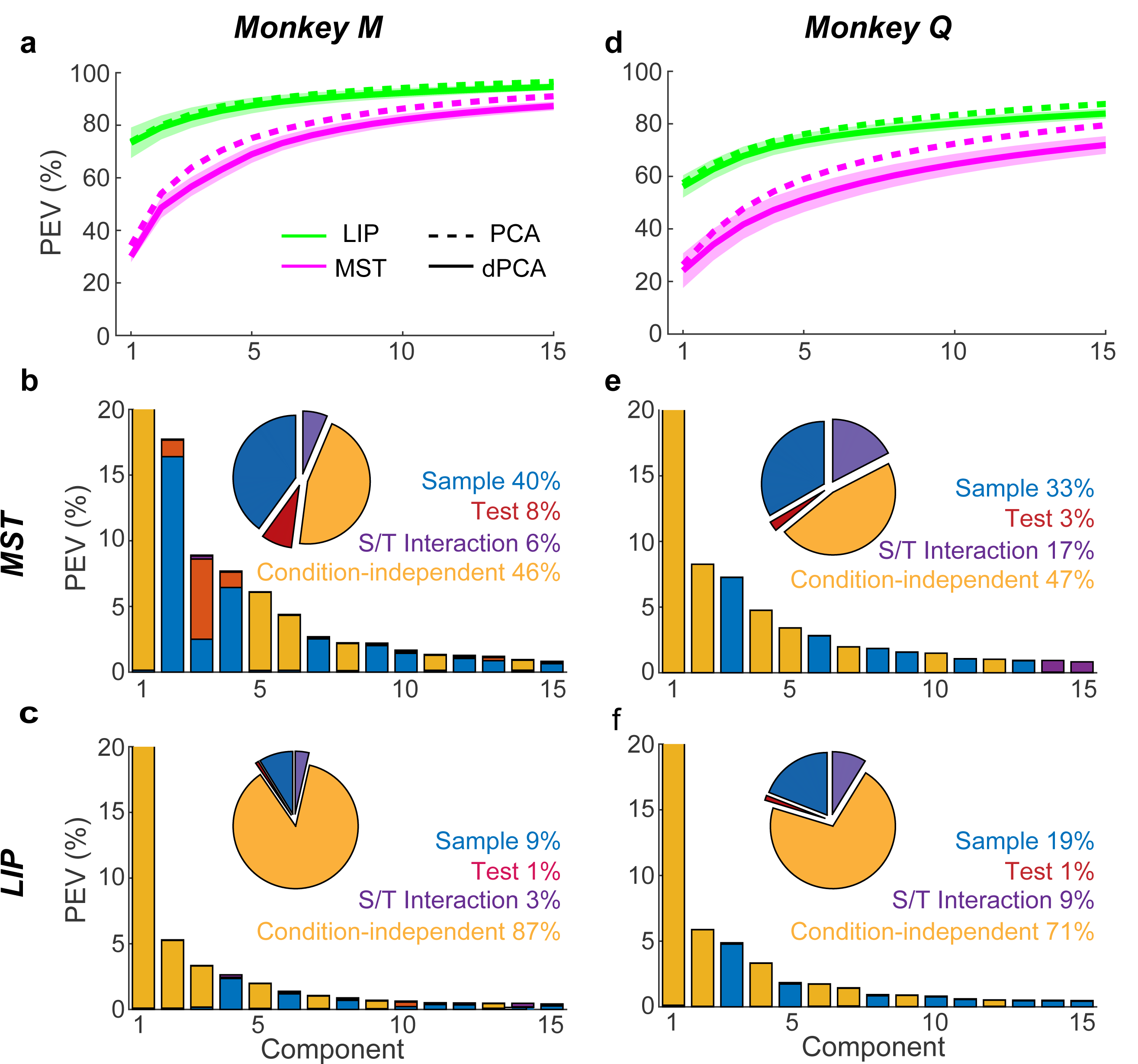
Demixed PCA (dPCA) applied to MST and LIP population activity in the DMC task. **a**. Cumulative variance explained by PCA (dashed) and dPCA (solid) for MST (pink) and LIP (green) population activity for monkey M. Only the first 15 principal components (PCs) were shown. dPCA explains almost the similar amount of variance as standard PCA (compare the solid line to dashed line). The shaded area denotes the ±STD of the dPCA. **b**. Variance of the individual demixed PCs of MST population activity for monkey M. Each bar shows the proportion of total explained variance, which is contributed by four different task components represented by different colors. The pie chart shows the total variance explained by each task variable. **c**. Variance of the individual demixed PCs of LIP population activity for monkey M. **d-f**. The results of dPCA applied to MST and LIP population activity of monkey Q.

### Decision-correlated neural activity in MST and LIP

To test whether sample category encoding in both MST and LIP was correlated with the monkeys’ trial-by-trial categorical decisions, we compared neuronal activity between correct and error trials. There are two potential scenarios for decision-correlated category selectivity, with each suggestive of a particular kind of correlation between neuronal category selectivity and monkeys’ decisions: 1) similar category selectivity on correct and error trials, indicating that stimulus tuning was fixed rather than varying with the monkeys’ decisions; 2) significantly different category selectivity between correct and error trials, suggesting that stimulus selectivity was closely and dynamically coupled to the monkeys’ decision process. Of particular interest is category selectivity that showed an opposite sign between correct and error trials, indicating a close relationship between neural category selectivity and the monkeys’ behavior. **Fig. 3a,b** show activity for example MST and LIP neurons on correct and error trials (same neurons as Fig. 2a,c). The example neurons from both areas showed significantly different category encoding between correct and error trials, and their category preferences were reversed in sign on error compared to correct trials during most of the task periods. This suggests that both MST and LIP activity was correlated with monkeys’ categorical decisions.

To quantify decision-correlated neural activity at the population level, we first trained sample category SVM classifiers, using activity on correct trials, and then tested their decoding performance on error trials. We only included trials in which the test motion directions were far (>= 30 degrees) from the boundary, as the errors on these trials were most likely due to mis-categorizing the sample stimulus. Sample category encoding which reversed its sign between correct and error trials would be expected to produce decoding performance values significantly below the chance level. Indeed, decoding performance dropped below chance shortly following sample onset and was maintained below chance throughout all task periods for both MST and LIP (**Fig. 3c,d**), indicating that population activity in both areas was significantly correlated with the monkeys’ trial-by-trial categorical decisions.

We examined stimulus-related and choice-related components of category selectivity for each single neuron from both areas using a partial correlation analysis. We calculated the r-stimulus (the partial correlation between neuronal activity and stimulus category, given the monkeys’ choices) and r-choice (the partial correlation between neuronal activity and monkeys’ categorical choice, given the stimulus category) for each neuron using both correct and error trials (see *Methods*). Only trials in which the sample directions were near the boundary but the test directions were far from the boundary were included in this analysis, as there were sufficient numbers of errors on these trials and these errors were most likely due to mis-categorizing the sample stimulus. This analysis revealed a significant correlation of neuronal activity with both the physical stimuli and monkeys’ categorical choices in all three periods of the DMC task and in both cortical areas (**Fig. 3e,f**). Meanwhile, neural activity in both MST and LIP was more closely correlated with monkeys’ categorical choices than the physical stimulus category during most of the task periods, indicated by the higher r-choice than r-stimulus values (p < 0.05, paired t test). Interestingly, LIP activity was more decision-correlated than MST during the early-to-mid sample period (r-choice, 150-350 ms after sample onset, p = 0.036, df = 799, tstat = -2.11, unpaired t-test), and became more choice-correlated than stimulus-correlated from the middle sample period (335 ms after sample onset). However, MST activity became more choice-correlated from the later sample period (615 ms after sample onset). Furthermore, LIP activity during the test period was more decision-correlated than MST activity (p = 0.0064, df = 799, tstat = -2.74, unpaired t-test).

Together, the decoding and partial correlation analyses indicate that sample category encoding was correlated with monkeys’ categorical decisions in both MST and LIP, with significantly stronger correlations observed in LIP than MST during the early sample period.

### Temporal stability of category encoding in MST and LIP

The differences between MST and LIP category encoding shown so far are likely to arise from the patterns of information flow between these areas for mediating the DMC task. To further test how sample category information transitioned into and was maintained during the delay period, we evaluated the stability of category encoding across all DMC task periods. Specifically, we aimed to test whether population category encoding was stable across different task periods by training and testing the category classifier using neural data from different time points in the trial. For example, if a classifier trained using neuronal data from one task period (e.g. sample period) showed high decoding performance when tested at another task period (e.g. delay period), this would indicate stable neuronal encoding between those task periods. **Fig. 4a,b** show the results from MST and LIP, respectively. In LIP, there were two stable periods of high category-classification accuracy spanning the sample and delay period respectively, indicated by the appearance of two separated rectangular regions of elevated classification accuracy during the sample and test periods. This indicates that the format of population category encoding in LIP was different between the sample and delay periods, and suggests a sharp transition of category encoding around the time of the early delay. In contrast to LIP, there were no obviously separated stable periods of category-classification for MST during the sample and delay periods, and the time period of high decoding performance for MST continued from the late sample into the early delay. This indicates that category encoding in MST was consistent between the late sample and early delay periods. Furthermore, a period of high-accuracy category classification was evident in LIP when training on the early sample and testing on the test period. However, this was not observed in MST. This indicates consistent sample-category encoding between sample and test periods in LIP. Together, these results suggest that LIP is more involved in encoding category information during the sample and test periods and maintaining category information in working memory,while MST appears more involved in transitioning category information from sensory encoding into working memory.

### Dimensionality of MST and LIP encoding

The results so far show that both MST and LIP activity encodes the visual motion categories, and that category encoding is decision-correlated in both areas. However, solving the DMC task requires a representation of multiple stimulus and task variables, including stimulus directions, sample and test categories, encoding sample information in working memory, and the comparison between sample and test stimuli. In order to assess the involvement of MST and LIP in representing multiple DMC task variables, we examined neural encoding of different task variables using a dimensionality reduction approach. We used demixed principal component analysis (dPCA) to decompose neural pseudopopulation activity into individual components which correspond to neural representations of different task variables^34^. Specifically, we decomposed population activity into four task-related variables: sample stimuli, test stimuli, sample-test interaction, and timing (condition-independent) (see *Methods*). This revealed two key differences between MST and LIP activity in the DMC task. First, dimensionality of the neural representations was greater in MST than LIP, as the first 10 principle components (PCs) explained much less variance of the population activity in MST (**Figure5 a,c,e,g,** shown separated by each monkey, MST vs. LIP: monkey M: 82.1±2.16% vs. 92.3± 1.82%, p < 0.001 monkey Q: 64.6± 3.93% vs. 80.1± 2.25%, p < 0.001, bootstrap). The greater dimensionality of MST population activity is also evident by directly estimating the dimensions of neural activity (p < 0.001 for each monkey, bootstrap, see methods). This suggest that neural activity in the DMC task is more homogenous across neurons in LIP than MST. The first PC of the population activity, which represents the stimulus-independent visual response to both sample and test stimuli in this case (**Extended Data Fig. 6**), explained more than 50% of the total variance of LIP population activity for both monkeys (monkey M = 73.3± 5.80 %, monkey Q = 56.2± 4.24 %), but less than 50% in MST (monkey M = 30.1±2.20%, monkey Q = 24.1± 6.57 %, p < 0.001 for both monkeys, bootstrap). The stimulus-independent visual response in PPC has been suggested to reflect visual salience and bottom-up attention^35,36^. Thus, this result is consistent with LIP showing a greater involvement in salience encoding than MST. Second, the task stimuli (sample, test, and sample-test interaction) explained much more variance of neural activity in MST than LIP (**Fig. 5 b,d,f,h**, p < 0.001 for each monkey, bootstrap), consistent with MST’s well-known role in visual motion processing.

**Fig. 6.**
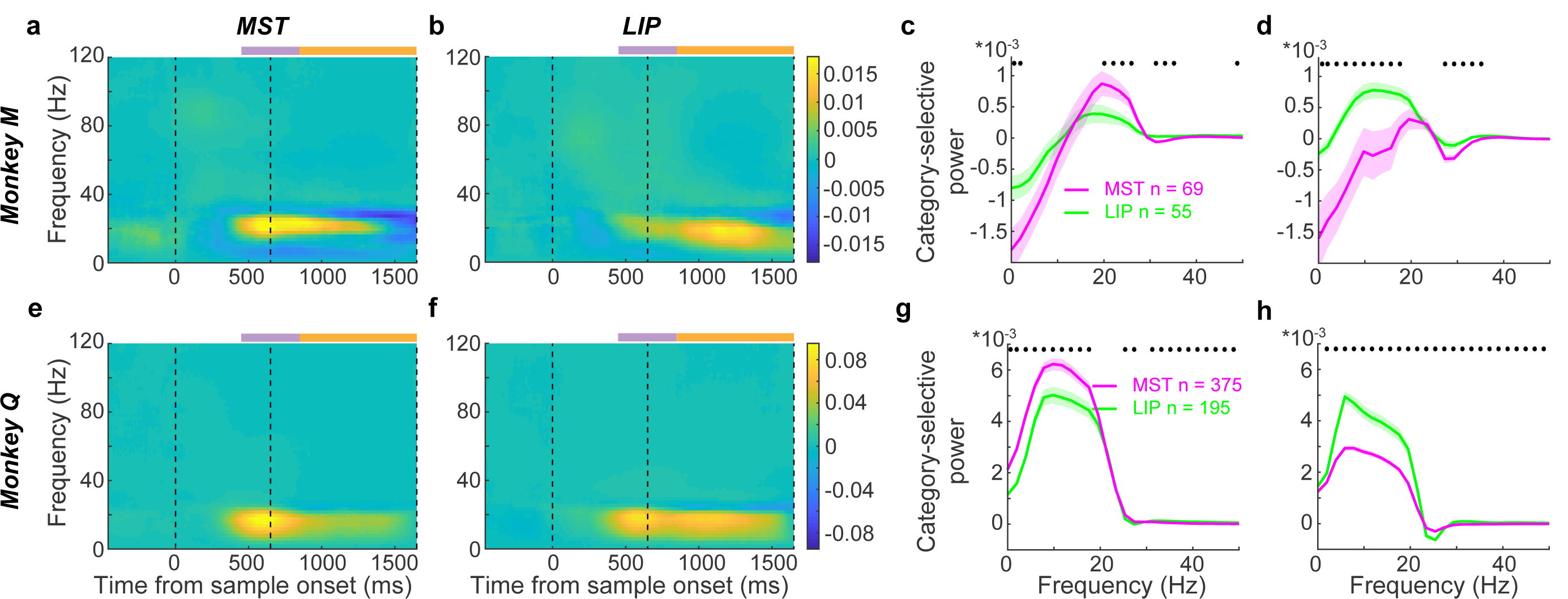
Beta-band oscillatory activity of LFPs in both MST and LIP encoded sample category information during working memory. **a-b**. Category-selective LFP activity for monkey M. **a**. The averaged sample category-selective LFP power recorded from MST electrodes in time-frequency space. The first and second dashed vertical lines denote the time interval for sample stimulus, while the third dashed vertical line indicates the time of test stimulus onset. Power is normalized to 1/f. The purple and orange bars mark the late-sample-to-early-delay as well as mid-to-late delay periods used in later analysis in e-h. **b**. Averaged sample category-selective LFP power recorded from LIP electrodes. Strong category-selective LFP activity is seen in the beta band during the mid-to-late delay period of the DMC task. **c-d**. The comparisons of category-selective LFP power between MST (pink) and LIP (green) during the late-sample to early-delay(e) and mid-to-late delay (f) periods for monkey M. The shaded area denotes ±SEM. The black star marks the frequency band for which there was significant difference between MST and LIP. **e-f**. Sample category-selective LFP activity in MST (c) and LIP (d) electrodes for monkey Q. **g-h**. The comparisons of category-selective LFP activity between MST and LIP for monkey Q.

### Decision and working memory encoding in LFP beta oscillations

Although previous studies of visual categorization primarily focused on analyzing category encoding in neuronal spiking activity, other studies have identified categorization-task related LFP activity^37,38^. Recent evidence suggests that LFP oscillations in PFC, particularly within the beta-band, might play a role in visual categorization. Beta-band synchrony within PFC, and between PFC and other brain areas such as anterior intraparietal cortex (AIP) and striatum, has been shown to represent task related information in categorization tasks^37,39^. However, the role of oscillatory activity in PPC during the visual categorization process has not been closely examined, except for one previous study which examined LFP oscillations in the anterior intraparietal (AIP) area during spatial categorization^37^. To better understand task-related LFPs in both MST and LIP during the DMC task, we computed the single-trial LFP power spectra for channels in which there was at least one task-related neuron, and then averaged them across trials based on the sample category (see *Methods*). We found significant oscillatory activity modulation during the DMC task both during different task epochs and by the different motion categories, particularly in the beta band (12-30Hz). The beta power of MST and LIP LFPs decreased during the sample and early delay periods, and then recovered to baseline during the mid-to-late delay (**Extended Data Fig. 7**). Interestingly, beta power was significantly modulated by sample category in both MST and LIP shortly before and during the delay in both monkeys (**Fig. 6a-d**), with most recording channels showing stronger beta power for one particular sample category (preferred) than the other sample category (non-preferred). The preferred sample categories were consistent between MST and LIP recording channels for both monkeys. Furthermore, the magnitude of beta modulation was different between MST and LIP during different task periods (**Fig. 6e-h**). MST showed significantly stronger category-selective beta activity than LIP during the late-sample-to-early-delay period (−200-200ms to sample offset, monkey M: 16-28Hz, p = 0.029, df = 122, tstat = - 2.20; monkey Q: 10-20Hz, p = 0.0096, df = 568, tstat = -2.60; unpaired t test), whereas LIP showed stronger category-selective beta activity during the mid-to-late delay period of the DMC task (201-1000ms after sample offset, monkey M: 10-22Hz, p = 0.026, df = 122, tstat = -2.26; monkey Q: 10-22HZ, p = 3.66e-10, df = 568, tstat = 6.38; unpaired t test).

**Fig. 7.**
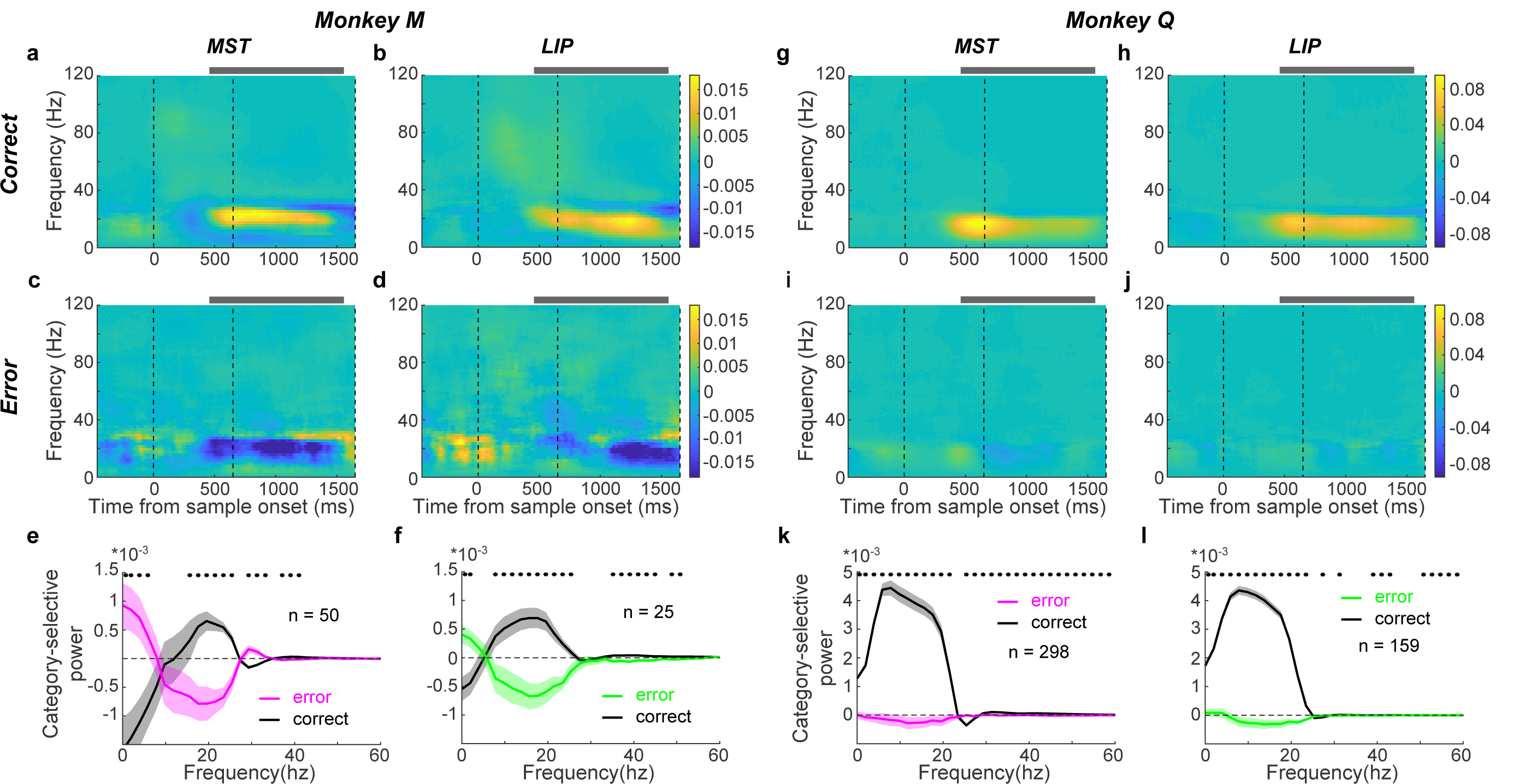
Beta-band LFP activity in both MST and LIP correlated with monkeys’ categorical decisions. **a-f**. Sample category-selective LFP activity on both correct and error trials for monkey M. **a-b**. Category-selective LFP activity in MST (a) and LIP (b) on correct trials. **c-d**. Category-selective LFP activity in MST (c) and LIP (d) on error trials. The grey bar marks the time period used for further analysis in e-f. **e-f**. Comparisons of category-selective LFP activity in MST (e) and LIP (f) between correct and error trials. The shaded area denotes ±SEM. The black star marks the frequency band for which there was significant difference between correct and error trials. **g-l**. Sample category-selective LFP activity on both correct and error trials for monkey Q.

To further test whether beta band LFP oscillations were correlated with monkeys’ categorical decisions, we compared category-selective beta activity between correct and error trials. As in the analysis of spiking activity, we only included trials in which the test directions were far from the category boundary (for which errors were most likely due to misclassifying the sample). As shown in **Fig. 7**, category-selective beta oscillations during error trials in both MST and LIP were either reversed in sign compared to correct trials (monkey M, **Fig. 7a-f**), or were greatly reduced in strength (monkey Q, **Fig. 7g-l**). These results indicate that the beta oscillations of LFPs in both MST and LIP were not merely a passive reflection of sample stimulus feature processing, but were instead correlated with working memory and decision processes.

## Discussion

We directly compared neuronal encoding between MST and LIP during a visual motion categorization task in an effort to understand how visual feature encoding is transformed into abstract categorical representations and decisions. First, we found that MST exhibits more flexible and task-related encoding than expected during the DMC task, with MST neurons showing abstract and decision-correlated encoding of visual motion categories with a similar strength as in LIP. This suggests that MST plays a greater role in cognitive functions than is widely assumed—going beyond its well-established role in visual motion processing. Second, we found substantial encoding of extraretinal task-relevant information in MST in both spiking and LFP neural activity during the working-memory delay period of the DMC task. This is consistent with a recent report of delay-period direction encoding during a motion-direction matching task, highlighting MST’s engagement with the frontal-parietal loop associated with working memory, and extends MST’s role toward more flexible or abstract decision making. Furthermore, LIP and MST differed in their time-courses of category selectivity, with greater category selectivity in the early sample period in LIP, whereas category selectivity in MST peaked in strength around the transition from the sample to delay period.

### Circuit mechanisms underlying motion categorization

Previous studies have given insight into the neural mechanisms underlying the transformation from visual feature encoding to more abstract categorical representations^1,2,40,41^. In particular, a line of studies from our group have focused on visual motion categorization across a hierarchy of visual, parietal, and frontal lobe cortical areas. We showed that neural activity in PFC, LIP, and MIP all encode learned categories during the DMC task, with evidence suggesting that LIP is more closely involved in the motion categorization process compared to MIP and PFC^3,8^. In contrast, neural activity in MT, an upstream motion processing area that provides input to LIP (and MST), showed strong direction encoding, but did not show an obvious encoding of the learned categories or altered direction tuning as a result of categorization training^1^.

In the present study, we show that neurons in MST, an important motion processing area within PPC, showed significant category encoding which was qualitatively similar to that in LIP during the DMC task. In addition, we compared neuronal category encoding between correct and error trials in both MST and LIP, and found such encoding in both areas was closely correlated with monkeys’ trial-by-trial decisions. These results are consistent with MST playing an important role in transforming upstream motion direction encoding (i.e. in MT) into abstract and task-related categorical representations. This indicates that multiple interconnected PPC subregions, including LIP, MST, and perhaps other PPC regions such as the ventral intraparietal area and 7a, are involved in the categorization process.

Although category encoding was broadly similar between LIP and MST, several pieces of evidence suggest that LIP might be more involved in the rapid categorization of visual stimuli than MST. First, category selectivity peaked in strength during the early sample period in LIP, but peaked in strength later in the sample period in MST. Second, category selectivity in LIP correlated more closely with monkeys’ trial by trial categorical decisions than in MST shortly following sample onset. These results are consistent with recent work from our group showing that LIP plays a causal role in motion-based categorical and perceptual decisions^11^. Category encoding in MST might arise from feedback from higher decision-related brain areas, such as LIP and PFC. Thus, it will be important for future studies to test whether MST also plays a causal role in the motion categorization process beyond its known role in visual motion processing.

It has been suggested that LIP plays a general role in encoding abstract or categorical information about visual stimuli^7,40,41^. A previous study showed that LIP neurons can encode both learned motion-categories and the learned pairings between associated shapes, suggesting a generality of category- or task-related encoding in LIP across multiple visual feature domains^7^. Meanwhile, studies from other groups have also shown that LIP represents a wide range of higher order factors beyond categorization, such as: task rules^42^, numerosity^43^, priority^44^, sensorimotor transformation^45^, motor error for corrective saccades^46^ and saccade timing^47^. Together, this evidence supports the view that LIP is generally involved in mediating abstract cognitive computations, beyond its role in visual-spatial functions such as attention and saccade planning. However, to our knowledge, MST has not been reported to encode visual features beyond visual motion, in addition to encoding vestibular stimuli and contributing to smooth pursuit eye movements^14,26,48,49^. Therefore, it will be important to test whether abstract encoding in MST is specialized for motion-based tasks, or whether it shows more generalized cognitive encoding during tasks based on other visual features as in LIP.

### Mnemonic encoding in MST and LIP and persistent activity

We observed that single-neuron activity and LFPs in both MST and LIP encoded categorical information during the delay period of the DMC task, and such mnemonic representations in both areas were correlated with the monkeys’ trial-by-trial decisions. This suggests that both areas are involved in maintaining category information in short-term working memory. However, delay period encoding differed between MST and LIP in several ways. First, delay-period category selectivity was stronger in MST than LIP during the early delay period, but this pattern was reversed at the end of the delay period (Fig. 2e) with LIP showing stronger late-delay category encoding. A similar pattern of results was evident in LFP activity, with stronger category-selective beta-band activity during the late sample and early delay periods in MST than LIP, but stronger beta-band encoding in LIP than MST during the mid-to-late delay. Meanwhile, the format of population category encoding in LIP differed between the sample and delay periods, as the category encoder trained in one period did not generalize to other time periods; whereas MST showed more consistent or temporally stable population category encoding throughout the late sample and early delay. Thus, MST activity appears more consistent with being involved in the transition of sample-period encoding into working memory, while LIP appears more involved in maintaining task-relevant information during the delay.

### Cognitive functions of MST

MST is reciprocally connected with several subregions of parietal cortex, including LIP, VIP and 7a^12^. MST is understood to be an important motion processing stage, but has been less implicated in higher cognitive functions. In particular, MST has been suggested to contribute to the perception of complex motion patterns, and integrating visual and vestibular signals for heading direction perception during self-motion^14,15,19,22,24-26^. During motion-based decision tasks, MST has been suggested to function as an intermediate processing stage between primary motion processing in MT and more cognitive processing in LIP^17,50^. Here we show that MST activity was closely correlated with monkeys’ categorical decisions in a similar manner as LIP, raising the possibility that MST is also causally involved in mediating such decisions. We also show that MST activity persistently encodes decision-correlated category information during the delay period of the DMC task, unlike upstream visual areas such as MT during the same task^1^. This is consistent with a previous study finding robust persistent direction-selective activity in MST in a delayed motion matching task^31^, but shows that delay-period encoding in MST extends to cognitive variables, beyond encoding of basic stimulus features. This also suggests that MST is more closely associated than MT with frontal-parietal circuits associated with working memory and task-related encoding. Previous work also found neural encoding in MST that appeared as intermediate between MT and LIP—for example, showing strong direction encoding as in MT with a modest influence of extraretinal or cognitive factors compared to that observed in LIP^29,30,32^. The more cognitive encoding observed in the present study could arise due to the specific demands of the categorization task compared to tasks used in previous MST studies. Another difference is in the method of neuron sampling (e.g. how neurons were selected and/or pre-screened before recording during the main task), as well as the areas of MST targeted for recordings. Most of the MST neurons in our study were recorded from MSTd, and we recorded from all task-related neurons which we encountered (see *Methods*), whereas previous studies appeared to more frequently sample motion direction selective neurons from lateral MST (MSTl). Therefore, it will be important for future studies to further test the differential roles of MSTd and MSTl in cognitive functions.

### Beta-band LFP activity during categorization and working memory

Frontoparietal oscillatory synchrony has been suggested to play a role in cognitive functions, such as attention^51^, visual working memory^52^, and decision-making^53,54^. Specifically, the beta oscillation (12-30HZ), often linked to motor functions, has been hypothesized to maintain the current sensorimotor or cognitive state via top-down selection of relevant neural ensembles^37,55^. There is increasing evidence that beta-band synchrony may also play a role in categorization and working memory. Previous studies have shown that beta-band LFP oscillatory coherence in both PFC and PPC is category-selective and may emphasize the encoding of task-relevant categories^37,38^. Interestingly, only category-selective PFC neurons, but not non-selective neurons, in those previous studies were synchronized with PPC beta oscillations, suggesting that long-range beta-band synchrony could act as a filter supporting task relevant encoding^37^. Category-selective beta-band synchrony has also been shown to develop between PFC and striatum in parallel with category learning^39^. Furthermore, beta-band synchrony between PFC and PPC as well as within PFC has been shown to encode stimulus information in working memory^52^. Consistent with previous studies, we found that beta-band LFP activity in both MST and LIP showed decision-related category representation shortly before and during the working memory delay, suggesting that PPC beta activity is involved in categorization and working memory. This is also consistent with the view that working memory is supported by frontal-parietal coordination. A previous study showed that beta-band LFP oscillations in PFC and AIP were especially prominent during the late sample and early delay periods of a rule based spatial categorization task^37^, whereas we found decreased beta-band LFP oscillations in LIP and MST during this period of the DMC task. Moreover, that study suggests that PFC circuits might compute the spatial categories and relay that information back to the parietal cortex, whereas our previous study showed that PPC is more likely to lead the motion categorization process compared to PFC^3^. These differences could be due to the different tasks (motion vs spatial categorization) and different PPC subregions (LIP, MST, and AIP) studied.

Overall, this study suggests that MST plays an non-trivial role in higher cognitive functions such as categorical decisions and working memory, extending beyond its traditionally recognized role in sensory processing of visual motion. It will be important to extend this work to examine whether MST plays a more general and causal role in cognitive functions beyond the tasks and stimuli tested here, as well as to examine the differential roles of the wider network of PPC subregions in flexible cognition.

## Acknowledgments

We thank Dr. Kenneth Latimer, Dr. Pantea Moghimi, Barbara Peysakhovich and Alessandra Silva for their constructive and helpful comments during the manuscript preparation.

## Author contributions

YZ and DJF designed the experiment. YZ trained monkeys and collected the behavioral and neuronal data, KM collected some of the neuronal data from LIP for monkey M. YZ analyzed the data and made figures. YZ wrote the manuscript. KM and DJF edited the manuscript. DJF supervised the experiments.

## Methods

### Behavioral task and stimulus display

The DMC task is similar as the tasks reported previously except that two near-boundary directions were added into each category. In this task, monkeys were trained to release a lever when the categories of sequentially presented sample and test stimuli matched, or hold the lever when the sample and test categories did not match. Stimuli consisted of 10 motion directions (15°, 35°, 55°, 75°, 135°, 195°,215°, 235°, 255°, 315°) grouped into two categories separated by a learned category boundary oriented at 45° (**Fig. 1b**). Trials were initiated by the monkey holding the lever and keeping central fixation. Monkeys needed to maintain fixation within a 2.5° radius of a fixation point through the trial. 500 ms after gaze fixation was maintained, a sample stimulus was presented for 650 ms, followed by a 1000 ms delay and a 650 ms test stimulus. If the categories of the sample and test stimuli matched, monkey needed to release a manual touch-bar within the test period to receive a juice reward. Otherwise, monkeys needed to hold the touch-bar during the test period and a second delay (150 ms) period, and wait for the second test stimulus, which was always a match, and then release the touch-bar. Therefore, monkeys concluded all trials with the same motor response (lever release). The motion stimuli were full contrast, 9° diameter, random-dot movies composed of 190 dots per frame that moved at 12°/s with 100% coherence. Task stimuli were displayed on a 21-inch color CRT monitor (1280*1024 resolution, 75 Hz refresh rate, 57 cm viewing distance). Identical stimuli, timing, and rewards were used for both monkeys in all LIP, and MST recordings. All ten motion directions were used for sample stimuli during all the recording sessions. Different from sample stimuli, ten motion directions were only used for test stimuli in about half of the recording sessions, while the four near-boundary directions were not used for test stimuli during the other half of the recording sessions. Monkeys’ eye positions were monitored by an EyeLink 1000 optical eye tracker (SR Research) at a sampling rate of 1 kHz and stored for offline analysis. Stimulus presentation, task events, rewards, and behavioral data acquisition were accomplished using an Intel-based PC equipped with MonkeyLogic software running in MATLAB (http://www.monkeylogic.net).

### Electrophysiological recording

Two male monkeys (Macaca mulatta, 8–12 kg) were implanted with a head post and recording chambers positioned over PPC. Stereotaxic coordinates for chamber placement were determined from magnetic resonance imaging (MRI) scans obtained before chamber implantation. We accessed area LIP and MST from the same PPC chamber, which was positioned over the intraparietal sulcus (IPS). Monkey M’s chamber was centered, under head stereotactic condition, at 10 mm lateral to the middle sagittal line, and 3 mm posterior to the middle coronal line, while Monkey Q’s chamber was positioned more laterally (centered ∼13mm lateral to the middle sagittal line, and 1.5 mm posterior to the middle coronal line) to gain more access to MSTd. Both chambers sit perpendicular to the sagittal plane. All experimental and surgical procedures were in accordance with the University of Chicago Animal Care and Use Committee and National Institutes of Health guidelines. Monkeys were housed in individual cages under a 12-h light/dark cycle. Behavioral training and experimental recordings were conducted during the light portion of the cycle.

LIP and MST recording sessions were interleaved in each monkey to reduce the influence of timing on the neuronal responses and monkeys’ behavior. In monkey M, 52 LIP recordings sessions were followed by 90 MST sessions and an additional 15 LIP sessions. For monkey Q, we first recorded 24 LIP recording sessions followed by 25 MST sessions, then conducted 8 LIP sessions followed by 21 MST sessions, and finally recorded 6 LIP sessions followed by 7 MST sessions.

The recording equipment and procedures were the same as in the previous studies^3,8,11^. All recordings on monkey M were conducted using single 75-μm tungsten microelectrodes (FHC). while most of the recordings on monkey Q were conducted using 16-channels (Plexon) linear v-probes after identified the locations of brains areas using single channel recording. In general, LIP neurons were found at more medial locations and MST neurons were found at more lateral locations within the same recording chamber. LIP was 4-8mm below the surface and MST was 4-10mm below the surface in both monkeys. Neurophysiological signals were amplified, digitized and stored for offline spike sorting (Plexon) to verify the quality and stability of neuronal isolations.

### Receptive field mapping and stimulus placement

Most LIP neurons as well as some MST neurons were tested with a memory-guided saccade (MGS) task before the DMC task. LIP neurons were identified by spatially selective visual or persistent activity during the MGS task for single channel recoding. During multi-channel recording, we included all the neurons recorded from the same grid locations and similar depths where we recorded spatially selective persistent activity neurons. MST neurons were identified by visual responses to visual motion patterns (whole screen expansion-contraction, as well as the linear motion stimuli used in the DMC task), and little or no modulation during the MGS task. For single channel recording in MST, we only recorded from neurons that showed activity modulation during the DMC task (but were not necessarily direction-selective). We included all the neurons recorded from the locations which were identified as MST for multi-channel recording. LIP and MST neurons were also differentiated based on anatomical criteria, such as the location of each electrode track relative to that estimated from the MRI scans, the pattern of gray– white matter transitions encountered on each electrode penetration, and the relative depths of each neuron.

The placement of motion stimuli differed slightly between single channel recording and multi-channel recording sessions. For single channel recording, motion stimuli for the DMC task were always placed in LIP neurons’ receptive fields (RFs), or the locations in the contralateral visual field which evoked the maximum visual response of MST neurons to the motion stimuli. The typical eccentricity of stimulus placement was ∼6.0-10.0°. For each multi-channel recording session, we first identified one stimulus or task responsive neuron according to above criteria, and then placed the motion stimuli according to the RF of the identified neuron.

### Data analysis

#### Pre-analysis neuron screening

We identified well-isolated singe units in both LIP and MST that showed task-related activity in the DMC task, using the following criteria: (1), the maximum averaged firing rate during at least one of the four different task periods (sample period, earlier delay period, later delay period, and test period) should be no less than 1 spike/s; (2), the activity should exhibit at least one kind of task-related modulation (such as: sample category selectivity, test category selectivity, sample direction selectivity, test direction selectivity, two-way nested ANOVA test, p < 0.01) during one of the four task periods, or the mean activity during at least one of the four task periods should be significantly different from the baseline activity (fixation period). In total, 326 LIP neurons and 571 MST neurons were included for further analysis.

#### ROC-based category tuning index (rCTI)

We used the rCTI measurement to quantify the category selectivity, which was described in detail in our previous work^8^ and defined as follows:

rCTI = BCD - WCD,

WCD = (2*| ROC(75,195) - 0.5| + | ROC(135,195) - 0.5|+ | ROC(75,135) - 0.5| + 2*| ROC(255,15) - 0.5| + | ROC(315,15) - 0.5| +| ROC(255,315) - 0.5|+2*| ROC(55,215) - 0.5| + | ROC(55,75) - 0.5| + | ROC(195,215) - 0.5| + 2*| ROC((35,235) - 0.5| + | ROC(35,15) - 0.5| +| ROC(255,235) - 0.5|)/16; BCD = (2*| ROC(75,15) - 0.5| + | ROC(75,315) - 0.5|+ | ROC(135,255) - 0.5|+ | ROC(135,15) - 0.5| + 2*| ROC(195,255) - 0.5| +| ROC(195,315) - 0.5|+2*| ROC(55,35) - 0.5| + | ROC(55,255) - 0.5| + | ROC(75,235) - 0.5|+2*| ROC(215,235) - 0.5| + | ROC(195,35) - 0.5|+ | ROC(215,15) - 0.5|)/16 ;

#### Identify category-selective neuron

We performed a shuffle analysis to determine whether a neuron showed significant category selectivity, by determining whether the rCTI value of this neuron was significantly above chance level. To obtain a null distribution (chance level), we shuffled the direction labels of trials within each session to calculate the rCTI, and bootstrapped for 500 times. The rCTI value was determined as being statistically significant if it was greater than 99% of values from the null distribution. We applied this method to the mean activity within each time bin during the task period spanning from 50 ms after sample onset to 200ms after test onset (300ms bin size, six time-bins in total). Neurons were identified as category-selective if their rCTI value was greater than the significant threshold in at least one time-bin.

#### Classification of “pure direction selective” neurons

We identified the pure direction selective neurons according to the following criteria: (1), the neuron should be not identified as category-selective by the above criteria; (2) there was significant difference between activity to the different motion directions (one-way ANOVA, p < 0.01).

#### Determine the latency of category selectivity

For each neuron, we defined the threshold of significant category selectivity based on rCTI value. We set the rCTI value, which was three times SD above the baseline rCTI values (calculated during the fixation period using 100ms bin size stepped by 5ms), as the threshold. The latency of category selectivity was defined as the middle time-point of the first time-bin at which the rCTI value exceeded the threshold for at least two consecutive time bins.

#### Support vector machine (SVM) decoding

Similar to previous studies^4,8^, we used linear SVM classifiers to separately decode sample direction and category from a pseudo-population from the two cortical areas. Activities from different neurons in one cortical area were treated as if they were recorded simultaneously although neurons were mostly recorded separately. In training the SVM classifier, a hyperplane that best separates the trials belonging to two (category classifier) or five (direction classifier) different classes was determined. Each class corresponds to one type of category identity for category classifiers and one motion direction for direction classifier. As in previous studies^4^, we wanted to eliminate the contribution of direction selectivity and category selectivity to the performances of the category classifiers and directions classifiers, respectively.

Therefore, to train and test the category classifier, we separated our trials into two groups based on the motion directions (see **Extended Data Fig**. 3a, bright and dark arrows). We trained one classifier using directions from one group and tested with the directions in the other group. A second classifier was constructed by switching the training and testing groups. We then averaged the performance of these two classifiers. In this case, we minimized the contribution of the direction selectivity into the performance of category classifier. For the direction classifier, we trained and tested two classifiers using the directions within each category separately, and then averaged the performances of the two classifiers. This eliminated the contribution of category selectivity to the decoding performance of the direction classifiers.

Decoding was applied to the mean firing rates of neurons within a 100 ms sliding window (10 ms step). For each neuron, we randomly selected 66% of trials to train the classifier and left the other 34% of trials for testing. We then randomly sampled, with replacement, 160 trials from the training set and 80 trials for the testing set for bootstrapping. For each iteration of the bootstrap, we randomly selected 100 neurons with replacement from the neuron population from each area to perform the analysis. In order to reduce the potential confound caused by uneven numbers of trials of different motion directions, a minimum number for trials of each motion direction was required for random sampling (10 and 5 trials for training and testing data of each direction, respectively). We bootstrapped all decoding analyses 200 times.

#### Partial correlation analysis

A partial correlation analysis was performed similar to previous studies^11^. For each trial during the DMC task, we obtained three parameters, i.e., the category identity of sample stimuli, the neuronal activity, and the monkeys’ categorical choice (deduced from monkeys’ behavior), for the calculation. The stimulus category was assigned with different values for different motion categories: 1 and -1 are used for category 1 and category 2, respectively. Different categorical choices were also coded as different values (1 for choosing category 1 and -1 for choosing category 2). Two measures were then calculated: r stimulus = r(neuronal activity, stimulus category| choice category), the partial correlation between neuronal activity and stimulus category, given the monkeys’ categorical choices; and r choice = r(neuronal activity, categorical choice | stimulus category), the partial correlation between neuronal activity and monkeys’ categorical choices, given the stimulus category. In **Fig. 3e,f**, the partial correlation analysis was applied on each neuron’s average firing rate in a 100ms window, advanced in 10 ms steps. To perform this analysis, we only included the trials in which the sample motion directions were close (10°) to boundary directions but the test directions were far from the boundary. This is because there were enough errors for these trials, and the errors on these trials were most likely due to mis-categorizing the sample stimulus which was close to the boundary.

#### Unbiased fraction of explained variance (FEV)

To test the correlation between category representation and the level of task difficulty, we quantified category selectivity for different direction pairs for each MST and LIP neuron. To quantify the amount of information that a neuron encoded about motion category that was independent of the absolute neuronal firing rate, we calculated the unbiased fraction of explained variance (FEV) in the neuron’s firing rate that could be attributed to sample category with the following:

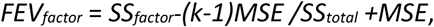

where *SS* indicates the sum of squares, MSE indicates mean square error, and k indicates number of conditions. In **Extended Data Fig**. 5, we first calculated the FEV values of the five opposite (180 degrees apart) motion direction pairs. Then, the FEVs of five direction pairs were combined into two groups according to their distance relative to the boundary (easier: ≥ 30°, more-difficult: 10°).

#### dPCA analysis

Demixed principal component analysis was performed using the methodology and code from a previous study (Kobak, Brendel et al.,http://github.com/machenslab/dPCA) that reduces the dimensionality of the population activity as the standard PCA and demixes all task variables. Specifically, we tested how much each task variable (sample stimuli, test stimuli, sample-test interaction, timing) contributes to the MST and LIP population activity during the DMC task.

As demonstrated in previous study^34^, the dPCA finds separate decoder (F) and encoder (D) matrices for each task variable (∅) by minimizing the loss function:

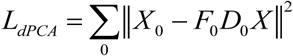

where X is a linear decomposition of the data matrix, which contains the instantaneous firing rate of the recorded neurons, into variable-specific averages:

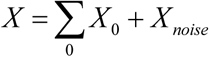

Here, we decomposed the neural activities into four parts: condition-independent, sample-dependent (10 sample directions), test-dependent (two test categories), dependent on the sample-test interaction, and noise. The decoder and encoder axes permit us to reduce the data into a few components capturing the majority of the variance of the data dependent on each task variable. In order to diminish the confound caused by the difference of population size between two brain areas to the dimension reduction analysis, we randomly sampled the population activity of 100 neurons with replacement from each brain area to perform the PCA and dPCA analysis. We then repeated the analysis 1000 times.

#### Estimate the dimensionality of population activity

Wecomputedthedimensionalityof the distribution of population vector responses in both LIP and MST, using the methodology described in a previous study^56^. These population vectors populate a cloud of points across many task conditions and time steps. The dimensionality is estimated as a weighted measure of the number of axes explored by that cloud:

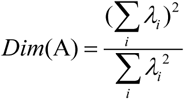

Where A denotes the population response matrix with each column representing one neuron, and λ_i_ is the i^th^ eigenvalue of the covariance matrix of A. The eigenvectors of the covariance matrix of A are the axes of the population response cloud. The resulted dimensionality can be thought as corresponding to the number of dimensions required to explain about 80% of the total variance.

#### LFP analysis

All the recording channels in which there was at least one task-related neuron recorded were included in the analysis in the current study. We excluded trials or sessions with artifacts in the LFP, such as those in which there were many time points (≥50ms) for which the LFP amplitude was clipped because of a mismatch in dynamic range between the amplifier and recording system. The LFP signal was pre-filtered by a band stop filter (Butterworth, 59-61 Hz) to remove power-line noise, and then z-scored in each recording session. We then used a MATLAB-based multi-taper analysis toolbox (chronux^57^) to analyze the power frequency spectra. The spectrograms were estimated using the LFPs within a 300 ms time window stepped by 10 ms.

In order to quantify the category selectivity in LFP activity, we first computed the single-trial power spectra of LFPs for each channel. We then averaged power spectra across trials based on the sample category, and calculated the difference of power spectra between two sample categories for each channel. We defined the sample categories for which most of the recording channels exhibited higher or lower beta power during the delay period as the preferred or non-preferred categories for each monkey, respectively.

