## Supplementary Figures for "Distributed Visual Category Processing Across Medial Superior Temporal and Lateral Intraparietal Cortices"

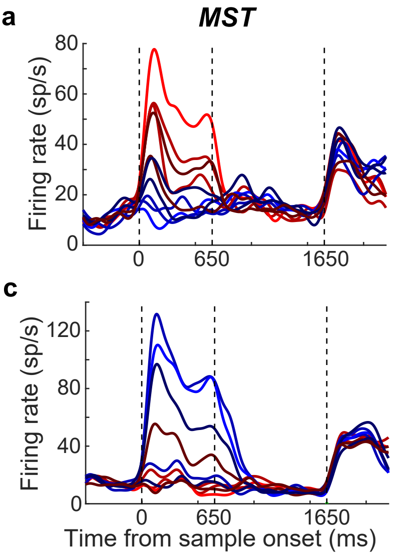


**Extended Data Fig. 1** Examples of pure motion direction selective neurons in MST and LIP. **a-b** Two example neurons from MST showed only direction selectivity in the DMC task. The averaged activity for each sample direction is plotted as a function of time. Different colors represent different sample categories, and different shades indicate the angular distance from the boundary. The ¬first and second dashed vertical lines denote the time for sample stimulus onset and offset, while the third dashed vertical line indicates the time of test stimulus onset. **c-d**. Two example neurons from LIP are shown in the same format as a-b.

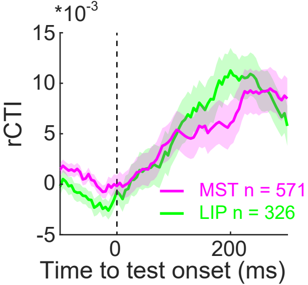


**Extended Data Fig. 2** Population level test category selectivity in MST (pink) and LIP (green) during the DMC task. A category tuning index (rCTI) shows the magnitude and time course of test category selectivity. Shaded area denotes ±SEM. The dashed vertical line indicates the time of test stimulus onset.

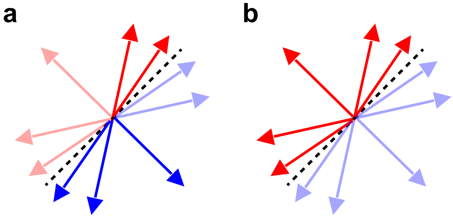


**Extended Data Fig. 3** Direction and category classification. **a.** Diagram depicting how motion category was classified independently of direction turning. As an example, neural responses to two directions from category 1 (bright red arrows) and three directions from category 2 (bright blue arrows) were used to train the classifier. The classifier was then used to test the category memberships of the remining five directions (dark arrows). A second category classifier was constructed by switching the training and testing directions, and the performance between these two classifiers was averaged. **b.** Diagram depicting how motion direction was classified independently of category selectivity. As an example, only neural responses to motion directions from category 1 (bright red arrows) were used to train and test the classifier. A second classifier was trained and tested using the directions from category 2, and the performance between these two classifiers was averaged.

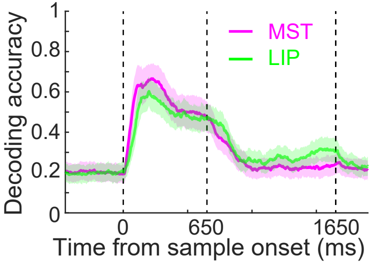


**Extended Data Fig. 4** Population level Sample direction selectivity in MST (pink) and LIP (green) in the DMC task. Time course of sample direction classification accuracy in MST (pink) and LIP (green) is measured using an SVM classifier. Shaded area denotes ±STD. The ¬first and second dashed vertical lines denote the time for sample stimulus onset and offset, while the third dashed vertical line indicates the time of test stimulus onset.

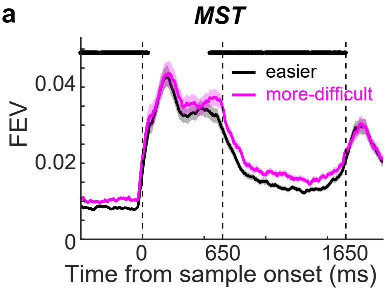


**Extended Data Fig. 5** The sample category selectivity in both MST and LIP correlated with the levels of task difficulty. The magnitude and time course of category selectivity for the easier and more-difficult trials was evaluated using fraction explained variance (FEV) for both MST (a) and LIP (b). The ¬first and second dashed vertical lines denote the time interval for sample stimulus onset and offset, while the third dashed vertical line indicates the time of test stimulus onset. The shaded area denotes ±SEM. The stars mark the time points for which there was significant difference between easier and more-difficult trials ($p < 0.01$, paired t test).

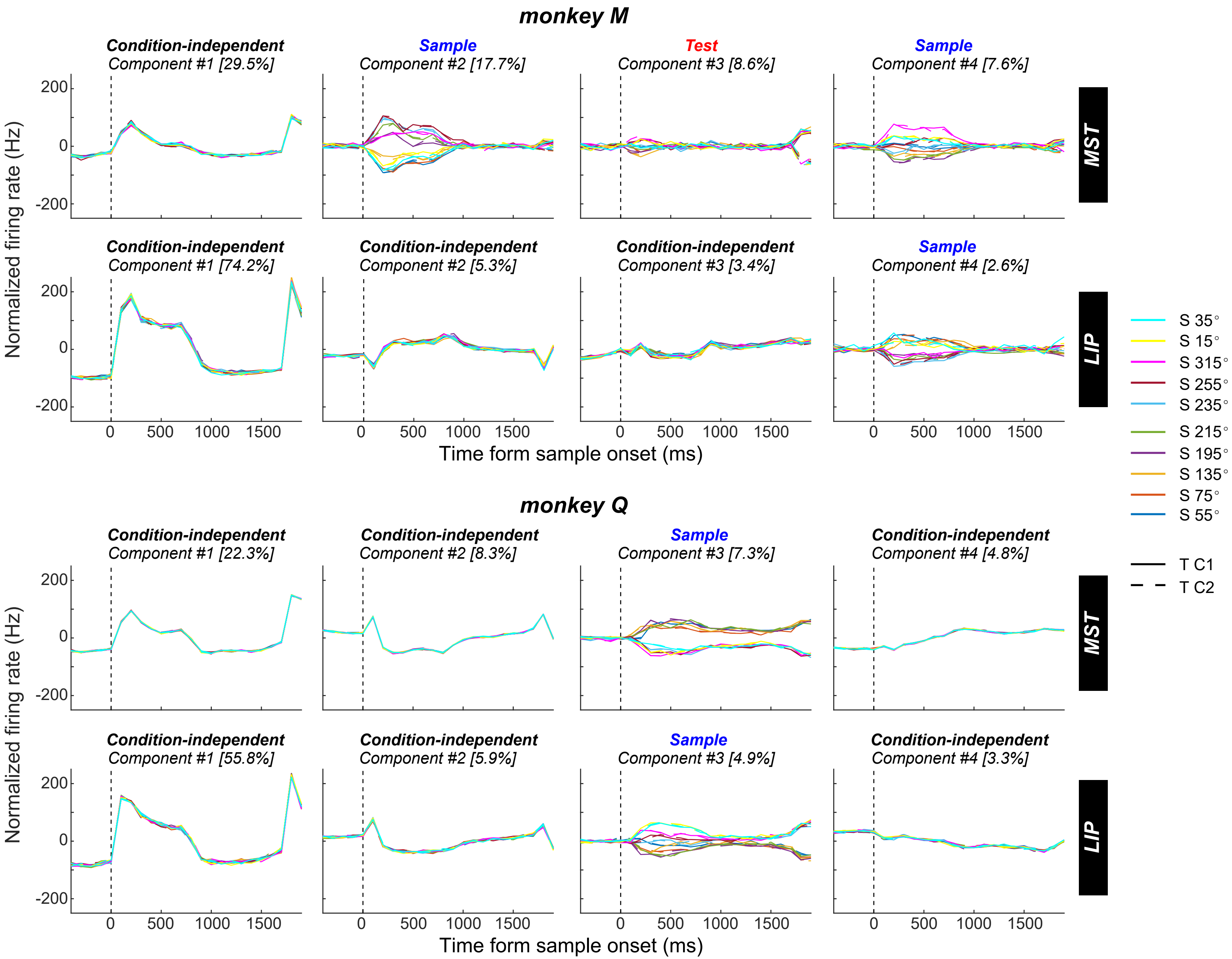


**Extended Data Fig. 6** Demixed PCs of MST and LIP population activity in the DMC task. The upper two rows: the results of dPCA applied to population activity for monkey M. Top row: the first four demixed PCs of MST population activity. In each subplot, the full data is projected onto the respective dPCA decoder axis, so that there are 20 lines corresponding to 20 conditions. Different colors represent different sample directions. Solid and dashed lines represent test category 1 and test category 2, respectively. The vertical dashed line marks the time of sample stimulus onset. Second row: first four demixed PCs of LIP population activity. The lower two rows: the results of dPCA applied to MST and LIP population activity for monkey Q.

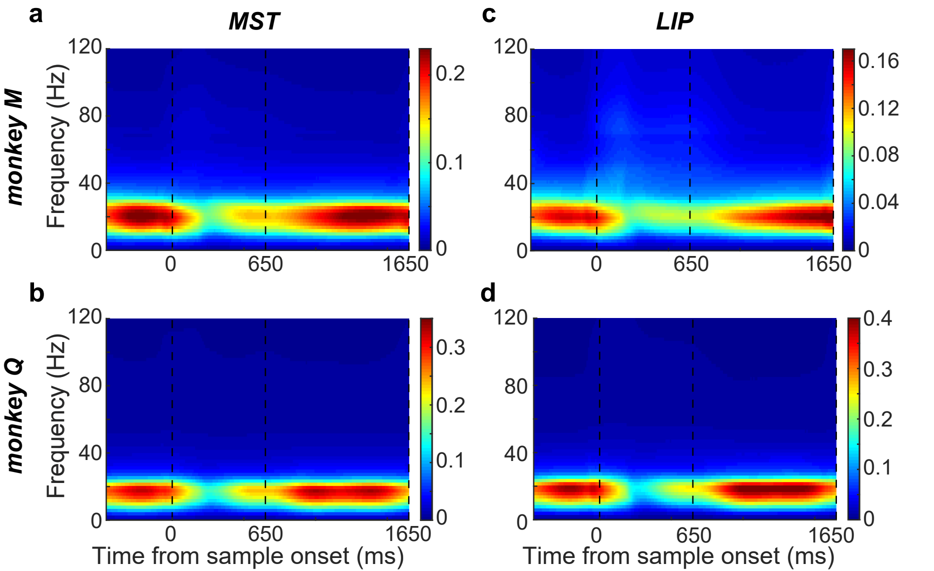


**Extended Data Fig. 7** The averaged LFP power recorded from MST and LIP during the DMC task. **a-b.** The averaged LFP activity recorded from MST (a) and LIP (b) for monkey M. a. The averaged LFP power recorded from MST electrodes in time-frequency space. The ¬first and second dashed vertical lines denote the time for sample stimulus onset and offset, while the third dashed vertical line indicates the time of test stimulus onset. Strong oscillation power was seen at the beta band (16–32 Hz) during the DMC task. The power is normalized to 1/f. b. The averaged LFP power recorded from LIP electrodes.  **c-d.** The LFP activity recorded from MST (c) and LIP (d) for monkey Q.
